# Differential representation of natural and manmade images in the human ventral visual stream

**DOI:** 10.1101/2022.09.22.509086

**Authors:** Mrugsen Nagsen Gopnarayan, Deeksha R. Rathore, Fabio S. Bauer, Jasper Hillard, Prerita Chawla

## Abstract

The human visual cortex processes visual stimuli hierarchically. Early visual areas (V1, V2) of the ventral visual stream feed crude visual features (like orientation and edges) into later visual areas (V4, lateral occipital (Lat Occ), inferior temporal (IT)) that then encode complex visual features (like object form). Previous studies have reported a difference in fMRI responses between natural and urban landscapes in certain parts of the brain. Here we asked if this distinction in the representation of complex natural and man-made visual stimuli extends to the ventral visual stream and if state-of-the-art convolutional neural networks (CNN) provide a map to this categorical distinction. To assess this, we used an open source fMRI data set of V1-4 and LatOcc BOLD responses to 1750 passively viewed natural and man-made grayscale images. The same images were fed into the pre-trained CORnet-S, a CNN designed to model hierarchical human visual processing layer-wise. To identify differences in representations within and between the human visual cortex and the CNN, we computed representational dissimilarity matrices. The BOLD response to the manmade and natural images shows correlation differences for the two categories, which increase across V4 and LatOcc. Such differences were also observed in the model CORnet-S for the layers V4 and IT. Our results suggest that the human visual cortex processes natural and manmade images differently starting from V4 and that this representational difference is modeled in CORnet-S. In both, categorical representation is progressively established in the later processing stages. This might indicate that the brain has perhaps developed two-distinct systems for their representations which can be directed through evolution. Further analysis may elucidate the contributory role of evolution towards this paradigm.

## 1 Introduction

Visual stimuli in humans are relayed through a series of downstream processing events, to the ventral and dorsal visual pathways that govern ‘what’ the stimulus looks like (object vision) and ‘where’ it is in space (spatial vision) (Ungerleider and Haxby [1994]). Both visual streams comprise the common V1 (primary visual cortex), V2, and V4 - collectively referred to as the early visual areas - where low-level image features (orientation, spatial frequency, color) are extracted from the incoming retinal signals of an image. However, at the level of V4, information branches into the dorsal and ventral streams, through neural projections to the posterior parietal and occipitotemporal regions respectively (Goodale and Milner [1992], Malach et al. [1995]). The lateral occipital complex (LatOcc) (which is homologous to the monkey inferior temporal (IT) cortex) (Orban et al. [2004]) and the IT cortex encode object shape and eventually object recognition via weighted coupling of low-level visual cues (Grill-Spector et al. [2001]). The ventral visual stream thus endows us with the remarkable ability to recognize objects in the outside world, everything from a tree or an animal to a bus or a building.

Several studies have reported differential fMRI blood oxygen-level dependent (BOLD) response activation to living (faces, body parts, animals) and non-living (places, tools, words) stimuli within the occipitotemporal cortex of the ventral stream (Kanwisher et al. [1997], Epstein and Kanwisher [1998], Downing et al. [2001], Martin et al. [1996]). This represents a category-specific modular representation of objects in the higher-level visual areas of the ventral stream. While this category speciation has been probed for individual object categories such as faces or tools or animals, the collective nature of ventral stream response involved in viewing natural vs man-made scene images is not clear. Man-made scenes (urban views, parks, bridges) have been a relatively recent addition to the long-standing backdrop of natural settings (forests, plains, mountains) over the course of evolution. Apart from the psychological and physiological benefits of viewing natural scenes that many studies have reported, it is also important to probe the nature of activation within the ventral stream to evaluate how broadscale settings are categorized within the visual hierarchy.

A study by Kim et al. showed that different brain regions are activated upon viewing natural and urban scenes. Within the occipital cortex, natural scenes activated the superior occipital gyrus in contrast to urban views which activated the middle and inferior occipital gyrus (Kim et al. [2010]). This directs to the possible role of early visual areas in the representation of the two classes. In this study, we probe the ventral stream for the visual areas involved in recognizing and distinguishing images of natural and manmade scenes. We also evaluate the performance of a convolutional neural network (CNN), CORnet-S, that closely mimics the human ventral visual stream (Kubilius et al. [2019]), in categorizing the two image classes, at each layer.

## 2 Methods

### Stimuli

Stimuli used for fMRI and CNN analysis were taken from the kay natural images data set. The dataset is publically available (link: https://crcns.org/data-sets/vc/vim-1/about-vim-1). All images were grey scale. The size of all the photographs was fixed at 20°×20° (500 pixels×500 pixels).

### Image labeling

Curating the dataset was essential for making sure the effect size we later found was genuine, and could not be explained by confounding factors by biases in the dataset. It was also critical to make sure that the distinction between “natural” and “manmade” was a genuine one that could have been quickly recognized by the human participants, but not a proxy for other unaccounted-for factors such as lighting. Our initial dataset was large enough to afford us some liberty in removing any images we suspected might elicit inappropriate responses in the subject’s visual cortices. Our researchers removed all scenes that could not quickly be determined as natural or manmade. Even if it “technically” was one or the other, our focus on neural processing could not be muddled by edge cases like a door in a landscape scene or unusual objects such as a starry sky. There were hundreds of such cases. Additionally, all images involving faces were removed. It is well known that the brain processes faces in a special way that is not of interest in this study and probably has no analog in the CNN. For other arousing stimuli, we removed them for similar reasons. Images were manually labeled using the image labeling feature of the site OpenAI (link: https://openai.com/about/). Each researcher on the team participated in removing images, examining each of them carefully to determine if they met the exclusion criteria. After the useful images were decided upon, we cross-checked some of the images selected by the other researchers to double-check that there were no inconsistencies. In this type of study, most types of bias in the selection of images would not hinder the results of the study, as the RDMs for analysis are highly extracted. Still, utmost care was taken in this stage to narrow the range of stimuli to the human brain into categories that we were interested in working with: natural and manmade.

### fMRI Dataset

The Kay natural images dataset (Kay et al. [2008]) was used to do the fMRI. It is publicly available at link: https://crcns.org/data-sets/vc/vim-1/about-vim-1. The dataset consists of BOLD fMRI responses to the passive viewing of image stimuli and the subjects performed no behavioral task while viewing images. The data collection was approved by the ethics committee of the Univesity of California, Berkeley. Two subjects S1 (age 33, M) and S2 (age 25, M) participated in the data collection. Both subjects had a normal corrected-to-normal vision. The bold responses for all the repeats of the images were averaged out and considered for analysis.

### Statistical analysis

The goal of this study is to highlight representational similarities between the ventral visual stream and an adequate model for two visual stimuli categories: man-made and natural scenes. A representational dissimilarity matrix (RDM) highlights correlational patterns out of a large set of stimuli-related activations. An RDM can aid comparisons of representational similarities between the brain and a model. We quantify this similarity of stimuli responses for manmade and natural stimuli by computing the representational dissimilarity matrix M for each cortical and each model layer. This matrix M is computed as one minus the correlation coefficients between z-scored population responses to each stimulus. The z-scored response of all neurons r to stimulus s is the response mean-subtracted across neurons i and normalized to standard deviation 1 across neurons i where N is the total number of neurons:

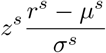

Where

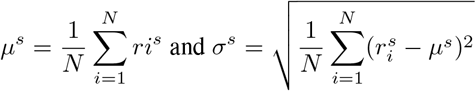

Then the full matrix can be computed as:

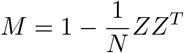

where Z is the z-scored response matrix with rows *r*^*s*^ and *N* is the number of visual stimuli of natural and manmade scenes (Jorge A. Menendez [2022]).

Since the RDMs are not easily readable and sufficiently informative, next we flattened the graphs to create histograms of Pearson r values for each ROI. We colored the histogram data differently depending on if it was brain or CNN data. For some of the layers, the differences between the distributions were visually apparent. For example, the V4 and Latocc regions assumed a distinctly bimodal shape, revealing a difference between natural and unnatural image encoding. After obtaining these histograms, we ran some statistical tests, first to verify the apparent normality of the RDM histograms. The Shapiro-Wilk test yielded test statistics of between 0.96 and 0.99 for each of the histograms. Despite the high test statistics, each of them showed statistically significant deviance from perfect normality, which is expected for large datasets. With the high test statistics and the visually apparent normality, we were satisfied to consider these datasets as being normal for the sake of comparing them.

## 3 Results

In this work, we wanted to assess if images of natural and manmade scenes are represented distinctly in the ventral visual pathway and if a representational difference in the human cortex is accurately modeled in a state-of-the-art CNN model of the human visual pathway.

The data BOLD fMRI data of V1, V2, V4, and LatOcc used for analysis was obtained from two individuals passively viewing 1870 grayscale images of 20×20px while fixating, each image is viewed 13 times (Kay et al. [2008]). The stimulus set was classified and subset to 785 natural and 345 manmade scenes for analysis. As a comparative model of the visual stream, we chose CORnet-S based on its high performance ranking in and its design with each model layer supposed to model the corresponding layer of the human visual cortex (Schrimpf et al. [2018], Kubilius et al. [2019], Schrimpf et al. [2020]). The features of the first four layers of CORnet-S, supposed to correspond to V1, V2, V4, and IT were used for analysis.

### 3.1 Visual stimuli of manmade and natural scenes are differently encoded in both the ventral visual stream and CORnet-S

To assess how similar or different the visual stimuli of manmade and natural scenes are being processed within the cortex and CNN, we computed the representational dissimilarity matrix (RDM). An RDM visualizes the similarity structure between representations of different stimuli. It can be concluded that a brain area and a model use a similar representational scheme if stimuli that are represented similarly in the brain are represented similarly in the model as well. Our analysis focuses on the early part of the visual stream. Thus with the underlying correlation coefficients of the z-scored stimuli responses computed, RDMs for each of the first four layers of the cortex and CNN were constructed. It is important to mention that the features extracted from the CORnet-S model are distinct from the z-scored fMRI BOLD responses computed and plotted. However, the model claims to resemble cortical response schemes well. The resulting stimuli axis on each RDM is sorted by category (natural, manmade) of stimulus to visualize any distinct representational schemes.

Figure 2 shows the RDMs of both fMRI and CORnet-S. For fMRI, the RDMs show hardly any strong distinction for any layer, especially when visually interpreting a low-resolution print of thousands of stimuli coming from a noisy source such as fMRI. Therefore the further quantitative analysis is needed. For the model, starting from the second layer a clear distinction between stimuli categories starts to form. This distinct representational scheme becomes increasingly apparent with each layer. This apparent representational scheme of the model and the strong difference in first layer dissimilarity between fMRI and model seem to point to room for possible improvements in CORnet-S.

**Figure 1:**
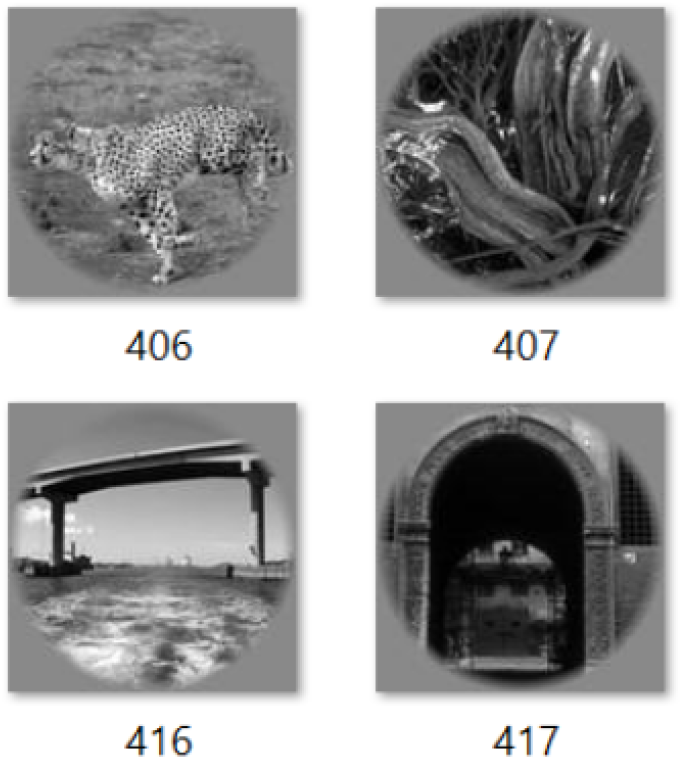
A sample of the 1750 passively fixated 20×20px grayscale images viewed 13 times each by the two participants in fMRI. The open source dataset was hand classified into 785 natural and 345 manmade images. Images that did not fit the two categories unambiguously or that showed full or partial human figures, were excluded.

**Figure 2:**
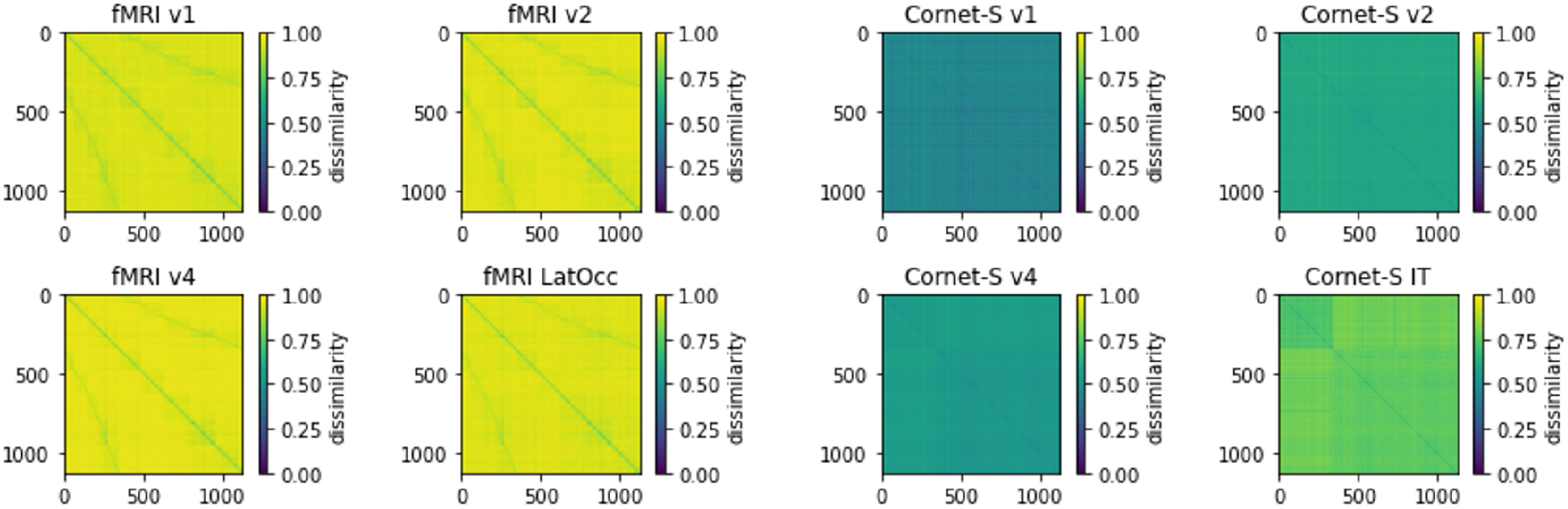
RDMs from the fMRI dataset and CORnet-S, representing a dissimilarity between natural and manmade images. The dissimilarity seems more pronunced and is easier visible in the model RDMs.

Thus the RDMs demonstrate a difference in the encoding of manmade and natural scenes in later layers of CORnet-S. In addition, the LatOcc activity of the human cortex starts to show signs of a distinct encoding as well. This first assessment needs to be quantified.

### 3.2 In both the CORnet-S and the visual cortex, representational differences become more significant in later stages of processing

What is the quantity of measured response dissimilarity for each layer and system? To quantify the measured response dissimilarity per layer and compare layerwise differences further, also across systems, we plotted the categorized response correlations for fMRI and model (figure 3) and computed Cohen’s distance for both (figure 4). The correlation differences were significant for all the layers in both Cornet and fMRI response (p < 1e-5, Man-Whitney test). For fMRI, the categorized responses to manmade and natural scenes start to show a difference in correlation in V4 and LatOcc. This apparent and quantitative correlational dissimilarity to the stimuli categories is pointing to a different representational scheme (Cohen’s d: V4 = 0.096, LatOcc = 0.191). For CORnet-S, representational schemes are similar to the fMRI data in that apparent as differences become apparent in V4 and are strongest in IT, the last layer analysis. As no fMRI data for IT was available, a direct comparison between the fourth model and the fMRI layer (LatOcc) is not possible. Nonetheless, a statement on similarities in the process can be made. As in the cortex, the model shows a sharp increase in dissimilarity between layers three and four (Cohen’s d: V4 = 0.176, IT = 1.037). The relative increase and absolute value of correlational differences are many times stronger in the idealized model, however.

**Figure 3:**
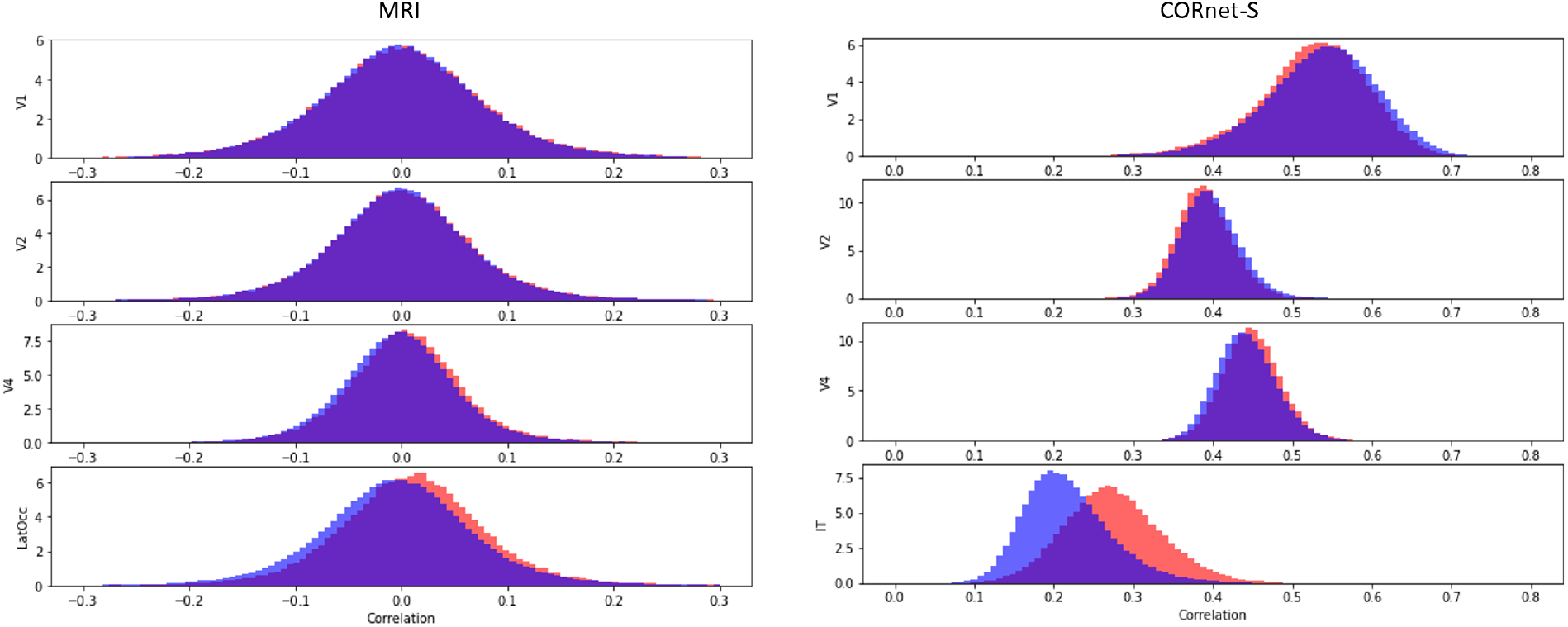
Histogram plots depicting the layerwise correlations obtained from the RDMs of fMRI dataset and CORnet-S. The difference in the correlation of natural and manmade images start appearing from the layer V4 and are strongest in LatOcc for human brain. In parallel, CORnet-S depicts the gradually increasing correlation differences for V4 and IT layer.

**Figure 4:**
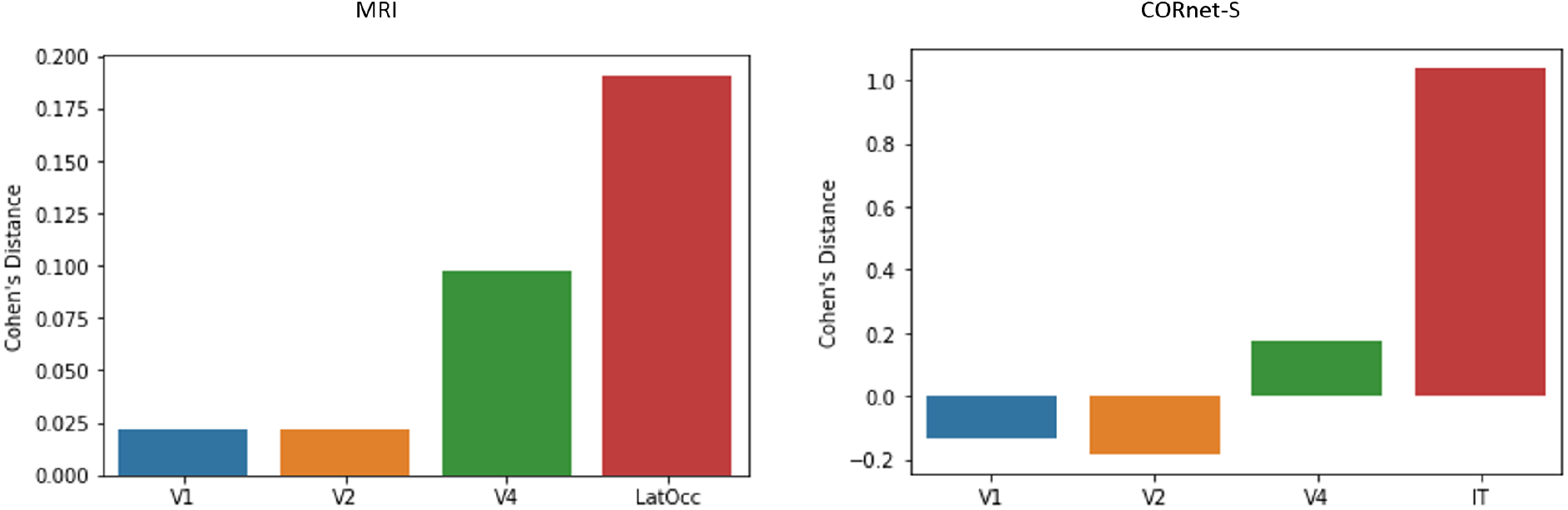
We calculated Cohen’s distance for each layer within both systems to be able to sufficiently judge the change and magnitude of representational dissimilarity between the model and brain. Both systems show an increase in the last analysed layer, with an apparent increase starting with the change from layer two to three. CORNet-S’s representational scheme for stimulus category results in a stronger distinction between natural and manmade images than in the brain. This might be due to the idealized model results, and a more noisy fMRI BOLD signal.

The analysis results suggest that images of natural and manmade scenes are represented distinctly in the early ventral visual pathway starting in V4, with a growing dissimilarity by layer, and that this representational scheme is modeled similarly in the CNN CORnet-S.

## 4 Discussion

For decades, the neural encoding of image stimuli has been an important problem in neuroscience. fMRI has proven to be the gold standard for studying neural encoding in humans due to its spatial resolution. The recent advances in the field of artificial neural networks have immensely helped us in analyzing the BOLD responses we obtain from fMRI scans. In our analysis, we tried to capture the power of decoding with fMRI scans to study the encoding of natural and man-made objects. Through our analysis, we could show that, indeed, there is differential encoding in the brain. But this is only limited to the regions V4 and Lateral Occipital (LatOcc), which are more downstream in the visual pathway. Earlier visual layers, V1 and V2, don’t show any differential encoding, which is consistent with our current knowledge about them being involved in lower-level pre-processing of the image stimuli. As the images in our dataset did not have a systematic difference at the lower feature level in terms of them being natural or man-made, there wasn’t a differential representation of images in layers V1 and V2 for being natural or man-made.

We also tried to check if these results also translate to the CORNet-S neural network, which is trained to mimic human object recognition. The layers V1, V2, V4, and IT in the Cornet-S are congruent to layers V1, V2, V4, and LatOcc in the human brain. As per our expectations, the results in CORnet-S were similar to the results we got for fMRI BOLD responses. The layers V1 and V2 in the CORnet-S showed similar representation, whereas the layers V4 and IT showed differential representation between natural and man-made images. The neural network layer IT, which is trained to mimic the brain region responsible for core object recognition, showed the most separation between the two image categories.

Our results in studying the representation in the brain were limited by the high level of noise in the BOLD responses of the fMRI responses. The use of better scanners and pre-processing techniques may lead to better neuro-imaging datasets, which would be more suited to answer the question about how exactly this differential representation takes place. The Kay natural images dataset also consisted of only grayscaled images, and as color is an important aspect of image recognition, a study using colored images may yield better insights. A neural network trained to classify natural and man-made images may help us decode the voxel level differences in image representation. Additionally, the fMRI dataset included only two subjects which limit the robustness of our analyses.

Overall, through our results, we can claim that there is a differential representation of natural and man-made images in the human brain, and this differential encoding is exclusive to only the latter regions in the ventral visual pathway. This differential encoding could be a result of either of multiple possibilities. One distinct module may have developed, over the course of evolution, to encode natural and man-made scenes separately in the V4 and layers thereon. Two, multiple modules responsible for encoding the contributory objects of a scene might be selectively active when viewing natural or man-made scenes. For example, the neural populations encoding a tree or an animal in a natural scene might be selectively activated to give rise to the wholesome representation of a natural scene and vice-versa for a man-made scene. Three, natural and man-made scene categorization could be a representation of their complex but distinctly different low-level image features themselves, suggesting a bottom-up processing event. A thorough analysis can be done by training ANNs first with natural images and then with man-made images to check for the possible role of evolution.

